# A generalization of t-SNE and UMAP to single-cell multimodal omics

**DOI:** 10.1101/2021.01.10.426098

**Authors:** Van Hoan Do, Stefan Canzar

## Abstract

Emerging single-cell technologies profile multiple types of molecules within individual cells. A fundamental step in the analysis of the produced high-dimensional data is their visualization using dimensionality reduction techniques such as t-SNE and UMAP. We introduce j-SNE and j-UMAP as their natural generalizations to the joint visualization of multimodal omics data. Our approach automatically learns the relative contribution of each modality to a concise representation of cellular identity that promotes discriminative features but suppresses noise. On eight datasets, j-SNE and j-UMAP produce unified embeddings that better agree with known cell types and that harmonize RNA and protein velocity landscapes. j-SNE and j-UMAP are available in the JVis Python package.

## Introduction

Single-cell RNA sequencing has enabled gene expression profiling at single cell resolution and provided novel opportunities to study cellular heterogeneity, cellular differentiation and development. Emerging single-cell technologies assay multiple modalities such as transcriptome, genome, epigenome, and proteome at the same time [3, 19, 23]. The joint analysis of multiple modalities has allowed to resolve subpopulations of cells at higher resolution [6, 9], has helped to infer the “acceleration” of RNA dynamics [7] and to extend time periods over which cell states can be predicted [18], and has linked dynamic changes in chromatin accessibiliy to transcription during cell-fate determination [1]. A fundamental step in the analysis of high dimensional single-cell data is their visualization in two dimensions. Arguably the most widely used nonlinear dimensionality reduction techniques are t-distributed stochastic neighbor embedding (t-SNE) [22] and uniform manifold approximation and projection (UMAP) [14]. Currently, these techniques are applied to each modality one at a time [3, 1, 4], and separate views of the data need to be reconciled by manual inspection. Here, we generalize t-SNE and UMAP to the joint visualization of multimodal single-cell measurements. While t-SNE and UMAP seek a low-dimensional embedding of cells that preserves similarities in the original (e.g. gene expression) space as well as possible, we propose j-SNE and j-UMAP that simultaneously preserve similarities across all modalities (Figure 1). Through Python package JVis they will allow to combine different views of the data into a unified embedding that can help to uncover previously hidden relationships among them. At the same time, our joint embedding schemes learn the relative importance of each modality from the data to reveal a concise representation of cellular identity.

**Figure 1:**
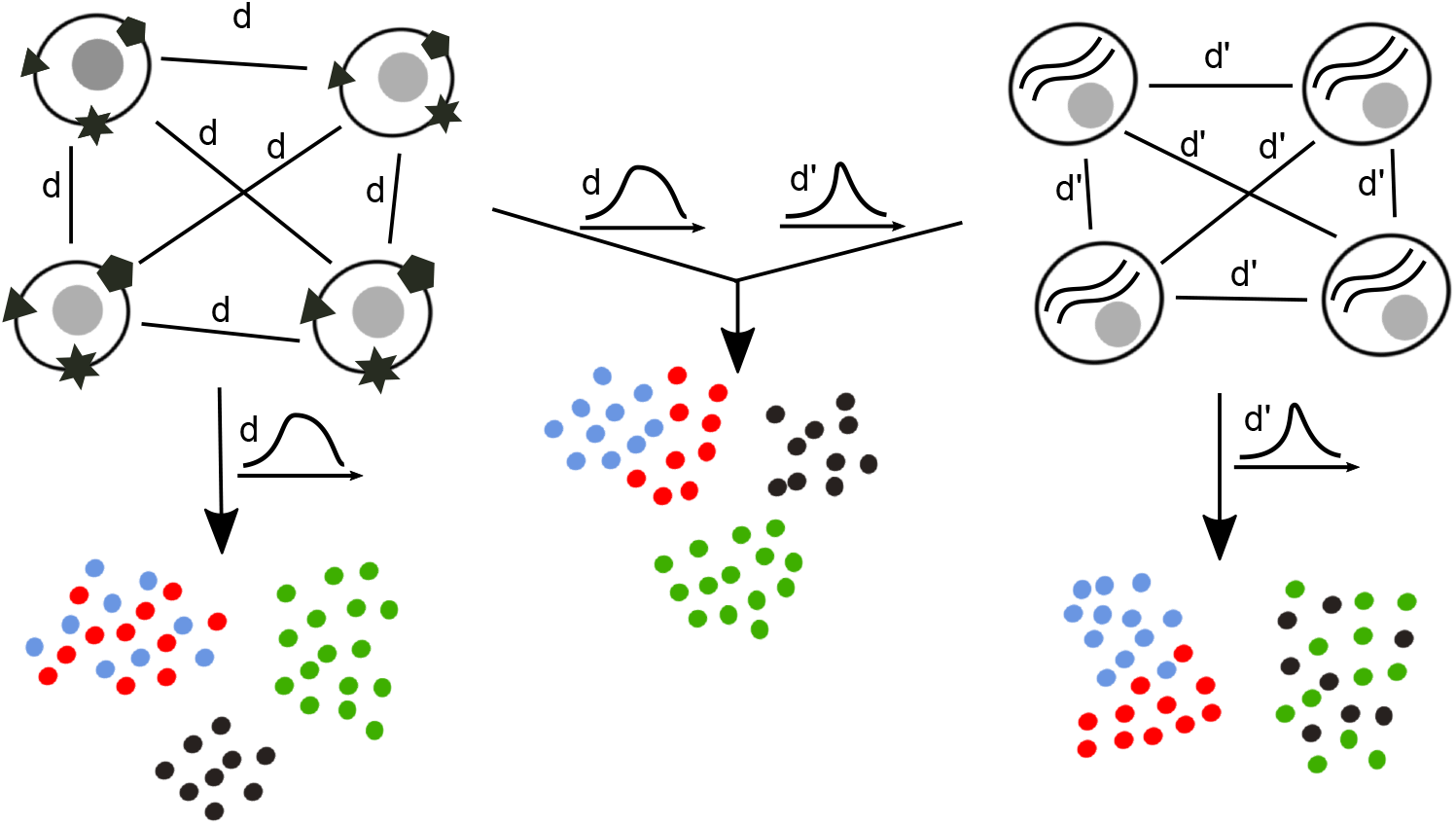
Overview of the joint embedding in JVis. Metrics *d* (left) and *d*′ (right) measure the dissimilarity of different cellular phenotypes of individual cells, such as the expression of surface proteins (left) and mRNA (right). t-SNE and UMAP learn a low-dimensional embedding of cells that preserves the distribution of similarities that are quantified based on *d* or *d*′ alone, which renders certain cell types indistinguishable to either modality. In this example, blue and red cells cannot be distinguished based on their measured surface proteins, and green and black cells overlap in transcriptomic space. In JVis we generalize t-SNE and UMAP to learn a joint embedding that preserves similarities in all modalities at the same time. We integrate *d* and *d*′ in a convex combination of KL divergences (j-SNE) or cross entropies (j-UMAP) between corresponding similarities in low and high-dimensional space. An arrangement of cells that minimizes this convex combination with simultaneously learned weights takes into account similarities and differences in both mRNA and surface protein expression to more accurately represent cellular identity (middle).

## Results

As proof of concept, we first demonstrate the ability of JVis to integrate modalities with different signal strengths. scRNA-seq, for example, often allows a finer mapping of cell states than singlecell ATAC-seq [20]. We used JVis to compute a joint embedding of accessible chromatin and gene expression measured simultaneously by SNARE-seq [4] in 1,047 single cells from cultured human cell lines BJ, H1, K562, and GM12878. Similar to the conventional t-SNE and UMAP embeddings of transcriptomes or chromatin state alone, our joint j-SNE and j-UMAP embeddings clearly separate cells into four distinct clusters (Supplementary Figures S1a-f). Even when randomly shuffling gene expression measurements between cell lines BJ and H1, JVis employs chromatin accessibility to disentangle mixed mRNA measurments and separate all four cell lines (Figures 2a-b and Supplementary Figures S1g-h). After adding a third modality that consists entirely of random noise sampled from a uniform distribution *U*[0, 100], the separation of the four cell lines persists in the j-SNE and j-UMAP embeddings (Figure 2c and Supplementary Figure S1i). Notably, JVis learns a weight for each modality from the data that reflects its relevance to the final embedding, with (unchanged) chromatin accessibility assigned the highest weight, partially perturbed gene expression obtaining a lower weight, and random noise assigned the lowest weight. JVis is thus able to automatically distinguish meaningful from noisy modalities to reveal the underlying structure of the data.

**Figure 2:**
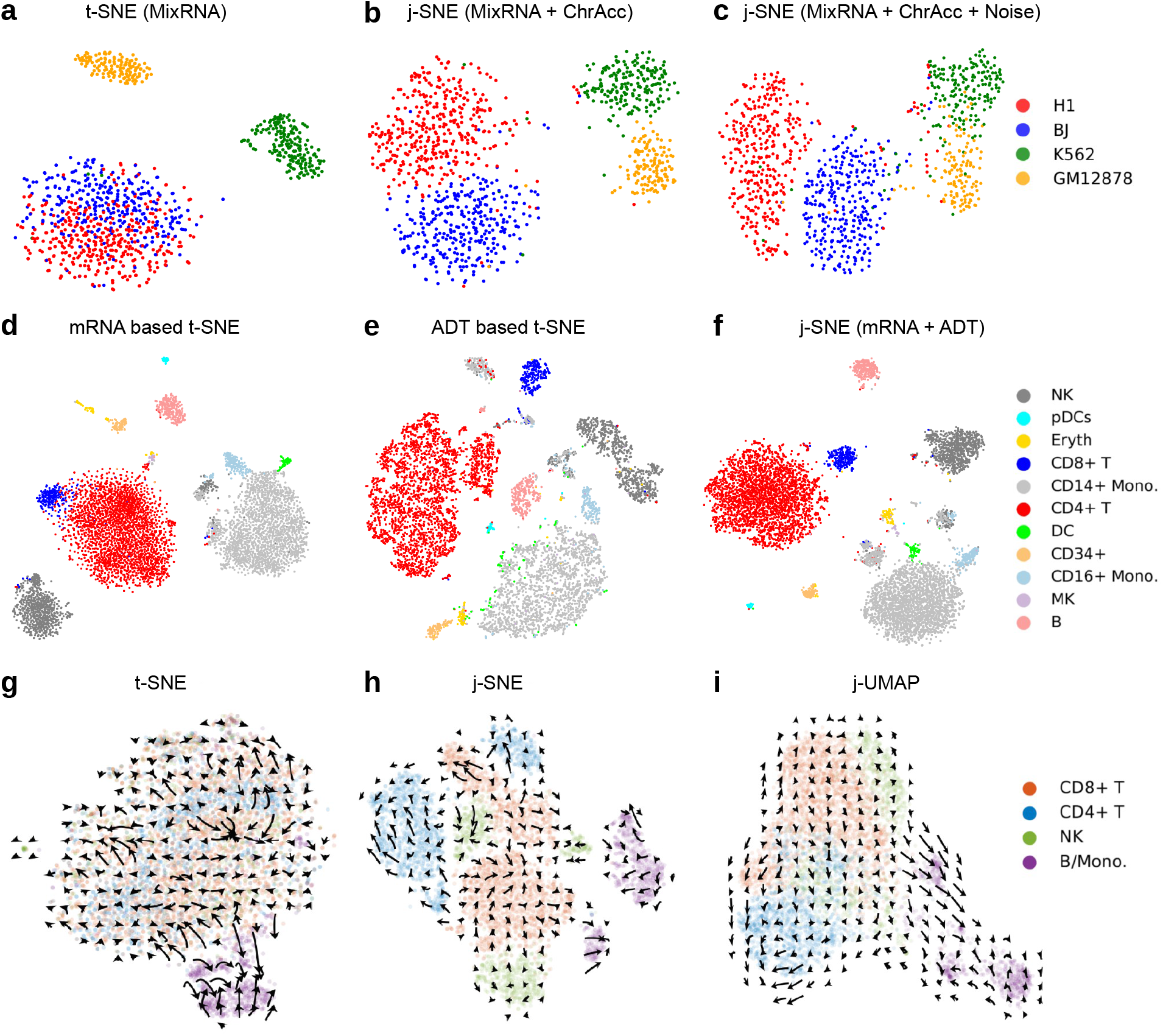
Comparison of cell types and protein acceleration in unimodal and multimodal embeddings. *First row*: Visualization of perturbed SNARE-seq measurements. Accessible chromatin (ChrAcc) and gene expression was measured simultaneously in single cell from human cell lines BJ, H1, K562, and GM12878. Gene expression measurements were randomly shuffled between cell lines BJ and H1 (MixRNA). (**a**) Conventional t-SNE embedding of cells based on shuffled gene expression alone. (**b**) j-SNE visualization of shuffled gene expression and (unchanged) chromatin accessibility. (**c**) j-SNE embedding of shuffled gene expression, chromatin accessibility, and a third modality that consist entirely of random noise sampled from uniform distribution *U*[0,100]. The simultaneously optimized weights assigned to the three modalities by JVis were *α* = (0.28,0.53,0.19), effectively suppressing noise. *Second row*: t-SNE/j-SNE visualizations of CBM cells. Cluster labels were identified by Specter. Embeddings were computed from RNA measurements alone (**d**), protein expression (ADT) alone (**e**), or jointly from both (**f**). *Third row*: Protein acceleration in ECCITE-seq (ctrl) data set projected into transcriptom-based t-SNE (**g**), and joint mRNA and surface protein based embeddings j-SNE (**h**) and j-UMAP (**i**).

In contrast, a naïve concatenation approach is not able to separate the perturbed measurements of cell lines BJ and H1 (Supplementary Figure S2). t-SNE and UMAP embeddings of concatenated vectors containing (mixed) RNA and chromatin counts cannot distinguish BJ from H1 cells and the addition of random noise as a third modality intermixes all 4 cell lines. While JVis borrows information across modalities and learns weights from the data that assign lower relative importance to the noise modality, the simple concatenation of modalities cannot benefit from their mutual dependence. To measure the accuracy of the embedding based on known cell types, we introduce metric KNI (see Supplementary Methods). KNI denotes the fraction on *k*-nearest neighbors in the embedding that are of the same type, averaged over all cells. A high KNI value indicates homogenous neighborhoods of cell types, while a random mixing of cells would cause low KNI values. In addition, we evaluate the quality of the embedding by measuring the agreement of clusters obtained by applying Louvain clustering [2] to the embedded cells with annotated cell types using the Adjusted Rand Index (ARI). In both experiments, JVis embeddings achieve higher KNI and ARI scores than conventional t-SNE and UMAP of concatenated modalities (Supplementary Table S1).

t-SNE and UMAP often produce embeddings that are in good agreement with known cell types or cell types computed by unsupervised clustering [2, 10] of high-dimensional molecular measurements such as mRNA expression. The simultaneous measurement of multiple types of molecules such as RNA and protein can refine cell types and JVis seeks to capture this refinement in their low-dimensional embedding. We compared unimodal and multimodal embeddings of mRNA and surface protein (ADT) expression measured in 4,292 healthy human peripheral blood mononuclear cells (PBMC) [15] and in 8,617 cord blood mononuclear cells (CBMC) [19] using CITE-seq [19]. Cell type labels were inferred by methods Specter [6] or CiteFuse [9], which have recently been introduced for the joint clustering of CITE-seq data.

Consistent with observations in [6, 9], t-SNE and UMAP visualizations of transcriptomic data alone does not show a clear distinction of CD4+ T cells and CD8+ T cells in the CBMC data set, while the embedding of protein expression mixes dendritic cells with CD14+ cells (Figures 2d-f, Supplementary Figures S3, S4). In contrast, JVis makes use of both modalities to compute a joint embedding that accurately separates CD4+ and CD8+ T cells as well as dendritic and CD14+ cells. Again, we confirm the visual interpretation quantitatively using the same metrics as above (Supplementary Table S2). The joint embedding of mRNA and ADT by JVis yields substantially larger ARI scores than the two unimodal t-SNE and UMAP emeddings. Similarly, the joint embeddings of cells in the PBMC data set by JVis separate naïve and memory CD4+ T cell that are mixed in the ADT based t-SNE and UMAP embeddings as well as CD4+ and CD8+ T cells that are mixed in the mRNA based embeddings (Supplementary Figures S5, S6). Again, joint embeddings yield more accurate clusterings in terms of ARI scores than unimodal embeddings (Supplementary Table S2).

RNA velocity [11] describes the rate of change of mRNA abundance estimated from the ratio of mature and pre-mRNA. While RNA velocity points to the future state of a cell, the recently introduced protein velocity [7] extends this concept and utilizes the joint measurment of RNA and protein abundance to infer the past, present, and future state of a cell. In [7], the authors used PCA and t-SNE to visualize RNA and protein velocity as well as the resulting *protein acceleration* in six PBMC data sets that were generated using four different technologies: CITE-seq, REAP-seq [17], ECCITE-seq [15] (data sets “CTCL”, a cutaneous T cell lymphoma patient, and “ctrl”, a healthy control), and 10X Genomics (data sets 1k and 10k). The authors observed strong velocity signals offered by the CITE-seq and 10x Genomics technologies, while REAP-seq and ECCITE-seq yielded noisier acceleration landscapes. Both RNA and protein velocity, however, were projected into the same t-SNE embedding of transcriptomic measurements alone, rendering their interpretation difficult. We therefore repeated the analysis of the six different data sets but projected velocities into the joint embedding of both modalities computed by JVis. The noisy acceleration landscapes observed in [7] in the ECCITE-seq and REAP-seq data sets become aligned across cell types in their joint embeddings by JVis (Figures 2g-i and Supplementary Figures S7, S8). Consistent with the improved distinction of transcriptionally similar CD4 and CD8 T cells in the joint embeddings above, acceleration landscapes in all six data sets are projected onto an embedding that more clearly separates CD4 and CD8 T cells compared to the original ones proposed in [7] (Figure 2g-i and Supplementary Figures S7-S11). RNA and protein velocities (without Bézier curve fitting for acceleration) for all six data sets are shown in Supplementary Figures S12-S17.

The noisy acceleration landscapes reported in [7] for the REAP-seq and ECCITE-seq data sets might be a result of the larger number of measured surface proteins (44 and 49 antibodies versus 13 and 17 antibodies in CITE-seq and 10X, respectively) that provide a finer distinction of subpopulations of cells. In fact, we observed lower agreement between RNA and protein based clusterings for the ECCITE-seq data set shown in Figure 2 (ARI 0.21), compared to the clusterings obtained from the two modalities in the CITE-seq data set that agree well (ARI 0.82). Since protein acceleration is computed from both RNA and protein abundances, their joint embedding can help to reduce visualization artifacts that arise when protein velocities are projected into a purely transcriptome based t-SNE embedding as in [7].

The complexity of Barnes-Hut based tSNE is 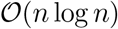, where *n* is the number of input cells [21]. Although no theoretical complexity bounds have been established for UMAP, its empirical complexity is 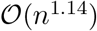 [14]. Therefore, JVis is expected to scale well to large data sets as it uses the same two underlying algorithms. For example, it took JVis less than 5 minutes to compute an embedding of the 10,000 cells contained in the largest data set used in this study (10x 10k). Moreover, our approach can easily be combined with the recently proposed FIt-SNE method [12] that scales linearly with the number of cells.

## Conclusions

t-SNE and UMAP are routinely used to explore high-dimensional measurements of single cells in low-dimensional space. We have introduced method JVis that generalizes t-SNE and UMAP to the joint visualization of single-cell multimodal omics data. We have demonstrated that JVis combines multiple omics measurements of single cells into a unified embedding that can reveal relationships among them that are not visible when applying conventional t-SNE or UMAP to each modality separately. Not surprisingly, projecting RNA and protein velocities into the joint embedding of both modalities yielded less noisy acceleration landscapes compare to embeddings of mRNA measurements alone. We therefore anticipate that JVis will aid in the meaningful visual interpretation of data generated by emerging multimodal omics technologies such as CITE-seq [19] and SHARE-seq [13], the latter allowing to combine RNA velocity with *chromatin potential*.

## Methods

A formal description of our generalizations j-SNE and j-UMAP as well as the algorithm to solve the underlying optimization problem can be found in Supplementary Methods. In brief, in j-SNE we want to learn a joint embedding of data points for each of which we have measured multiple modalities. Analog to t-SNE [22], we want to arrange points (here cells) in low-dimensional space such that similarities observed between points in high-dimensional space are preserved, but in all modalities at the same time. Generalizing the objective of t-SNE, we aim to minimize the convex combination of KL divergences of similarities in the original high-dimensional and similarities in the embedding low-dimensional space for each modality, where coefficients *α* of the convex combination represent the importance of individual modalities towards the final location of points in the embedding. We add a regularization term that prevents the joint embedding from being biased towards individual modalities. In j-UMAP we generalize UMAP to multimodal data analogously, minimizing a convex combination of cross entropies instead of KL divergences. In all experiments, we set the regularization parameter λ to 3 for j-SNE and to 1 for j-UMAP. We jointly optimize the location of points in the embedding and the importance coefficients *α* of modalities through an alternating optimization scheme: We fix coefficients *α* and find the best point locations by gradient descent, and in turn find optimal coefficients *α* for fixed locations by solving a convex optimization problem.

We measure the accuracy of an embedding using two different metrics. We introduce the *k*-nearest neighbor index (KNI), which denotes the fraction of k-nearest neighbors in the embedding that are of the same type. We used *k* = 10 and computed the average across all points. In addition, we use the agreement between true cell labels and labels obtained by a clustering of the embedded points as proxy for the quality of the embedding. The agreement between the two cell labelings is measured by the Adjusted Rand Index (ARI) [8].

Unless specified otherwise, clusterings of cells in this study are computed using the Louvain algorithm [2] where the resolution parameter is tuned to match the number of annotated cell types. We used standard preprocessing of the input data including log-transformation of the expression matrix followed by principal component analysis (PCA) for dimensionality reduction to 20 principle components.

We computed protein acceleration using the protaccel Python package introduced in [7].

## Declarations

### Availability of data and materials

The SNARE-seq and CBMC CITE-seq data sets were downloaded from Gene Expression Omnibus with accession codes GSE126074 and GSE126310, respectively. The six data sets used in the protein acceleration experiments were from [7]. The implementations of j-SNE and j-UMAP are based on the *scikit-learn* v0.23.1 library [16] and the UMAP v0.4.5 Python package [14], respectively. The JVis Python package can be installed through PyPi [5] and its open-source code is maintained at https://github.com/canzarlab/JVis-learn. Python scripts to reproduce all results in this paper are available at https://github.com/canzarlab/JVis_paper.

### Author’s contributions

V.H.D. and S.C. conceived the algorithm. V.H.D. implemented the software and performed the computational experiments. S.C guided the research. All authors wrote the manuscript.

