## Supplementary Methods, Tables, and Figures for "A generalization of t-SNE and UMAP to single-cell multimodal omics"

#### Generalizing t-SNE to multimodal data in j-SNE

Given $n$ data points $\mathcal{X} = \{x_1, x_2, \ldots, x_n\}$, t-SNE is a nonlinear dimensionality reduction technique that aims to learn an embedding $\mathcal{E} = \{y_1, y_2, \ldots, y_n\}$ in a low-dimensional space that preserves the distribution of point similarities. Based on a given metric $d$ that measures the dissimilarity between pairs of points, t-SNE first computes joint probabilities $p_{ij}$ that quantify the similarity between $x_i$ and $x_j$:

$$p_{j|i} = \frac{\exp(-d(x_i, x_j)^2/2\sigma_i^2)}{\sum_{k\neq i} \exp(-d(x_i, x_k)^2/2\sigma_i^2)}, \qquad p_{i|i} = 0, \tag{1}$$

$$p_{ij} = \frac{p_{j|i} + p_{i|j}}{2n}, \tag{2}$$

where $\sigma_i$ is the bandwidth of the Gaussion kernels centered at data point $x_i$. Given a predefined *perplexity* $u$, bandwidths $\sigma_i$ are chosen such that the perplexity of conditional distributions $P_i$ equal to $u$. The perplexity is often set to a value in the range between 5 and 50 [1]. t-SNE uses the normalized Student-t kernel with a single degree of freedom to measure similarities $q_{ij}$ in the embedding $\mathcal{E}$:

$$q_{ij} = \frac{(1 + \|y_i - y_j\|^2)^{-1}}{\sum_{k\neq i}(1 + \|y_i - y_k\|^2)^{-1}}, \qquad q_{ii} = 0. \tag{3}$$

To determine the location of points in the embedding $\mathcal{E}$ that preserve the original similarities $p_{ij}$ as well as possible, t-SNE seeks to minimize the Kullback-Leibler (KL) divergence of distribution $P = (p_{ij})$ from distribution $Q = (q_{ij})$.

$$C(\mathcal{E}) = KL(P||Q) = \sum_{i\neq j} p_{ij} \log \frac{p_{ij}}{q_{ij}}. \tag{4}$$

The minimization of equation (4) is performed using a gradient descent algorithm. The gradient is given by

$$\frac{\partial C}{\partial y_i} = 4 \sum_{j\neq i} (p_{ij} - q_{ij}) q_{ij} Z (y_i - y_j), \tag{5}$$

where $Z = \sum_{k\neq l}(1 + \|y_k - y_l\|^2)^{-1}$. While a naive implementation of t-SNE has runtime complexity $\mathcal{O}(n^2)$, the algorithm proposed in [2] uses the Barnes-Hut algorithm [3] and a sparse approximation of point similarities to reduce its complexity to $\mathcal{O}(n \log n)$ which allows it to scale to data sets comprising hundreds of thousands of cells.

We generalize t-SNE to the joint dimensionality reduction of multimodal data in j-SNE as follows. Given $K$-modal data points $X = \{x_1^{(k)}, x_2^{(k)}, \ldots, x_n^{(k)}\}, k = 1, 2, \ldots, K$, we want to learn a joint embedding $\mathcal{E} = \{y_1, y_2, \ldots, y_n\}$ that preserves similarities between points in $X$, analog to t-SNE, but in all modalities at the same time. Here, joint probability $p_{ij}^k$ measures the similarity of points $x_i^k$ and $x_j^k$ with respect to modality $k$. Generalizing the objective of t-SNE, in j-SNE we aim to minimize the KL divergence of distributions $P^{(k)}$ for each modality $k$ from distribution $Q$:

$$C(\mathcal{E}) = \sum_k \alpha_k KL(P^{(k)}||Q) + \lambda \sum_k \alpha_k \log \alpha_k \tag{6}$$

$$= \sum_k \sum_{i \neq j} \alpha_k p_{ij}^{(k)} \log \frac{p_{ij}^{(k)}}{q_{ij}} + \lambda \sum_k \alpha_k \log \alpha_k, \tag{7}$$

where $\alpha_k \geq 0$, $\sum_k \alpha_k = 1$, and $\lambda > 0$ is a regularization parameter. The first term of (6) denotes a convex combination of $\mathrm{KL}(P^{(k)}||Q), k = 1, 2, \ldots, K$, in which coefficients $\alpha_k$ represents the importance (contribution) of modality $k$ to the joint embedding. That is, the larger $\alpha_k$, the stronger the influence of modality $k$ on the final location of points in the embedding. The second term of the objective function is a regularization term that prevents the joint embedding from being biased towards individual modalities.

##### Optimization algorithm

We use an alternating optimization scheme to minimize (7) jointly over the location of points $y_i$ and the weighting of modalities $\alpha$. In each iteration, we first fix $\alpha$ and use t-SNE to find points $y$ in the joint embedding, after which we find the best (according to (7)) weighting $\alpha$ for fixed points $y$.

**Initialization of $\alpha$.** We initially give uniform weights $\alpha$ to all modalities, i.e.,

$$\alpha_k = 1/K, \text{ for all } k = 1, 2, \ldots, K. \tag{8}$$

**Step 1: Fix $\alpha$ and optimize over $y$.** Similar to conventional t-SNE, in j-SNE we employ the gradient descent algorithm to minimize (7). The gradient is given by:

$$\frac{\partial C}{\partial y_i} = 4 \sum_k \sum_{j \neq i} \alpha_k (p_{ij}^{(k)} - q_{ij}) q_{ij} Z (y_i - y_j) = 4 \sum_{j \neq i} \Big( \sum_k \alpha_k p_{ij}^{(k)} - q_{ij} \Big) q_{ij} Z (y_i - y_j). \tag{9}$$

Comparing gradient (9) to the original gradient (5), the former can be obtained by replacing distribution $P$ used in (5) by the convex combination of distributions $P^{(k)}$, i.e., $P = \sum_k \alpha_k P^{(k)}$. In other words j-SNE, the joint t-SNE of multiple modalities, can be computed by applying conventional (unimodal) t-SNE to the convex combination of distributions $P^{(k)}$ for all modalities $k$.

**Step 2: Fix $y$ and optimize over $\alpha$.** Given a joint embedding of points $y$, problem (7) is a special case of the following problem:

$$\begin{aligned} \min_{w} \quad & a_i\alpha_i + \lambda \sum_k \alpha_k \log \alpha_k \\ \text{subject to} \quad & \sum_k \alpha_k = 1, \\ & \alpha_k \geq 0, \qquad k = 1, 2, \ldots, K. \end{aligned} \tag{10}$$

This problem has a convex objective and linear constraints. We derive a closed form solution using the Karush-Kuhn-Tucker (KKT) conditions. The Lagrangian function corresponding to the constrained optimization problem (10) is given by

$$\mathcal{L}(\alpha, u, v) = a_i\alpha_i + \lambda \sum_k \alpha_k \log \alpha_k + u\big(\sum_k \alpha_k - 1\big) - \sum_k v_k \alpha_k.$$

The KKT conditions are given by

$$\begin{cases} \frac{\partial \mathcal{L}(\alpha,u,v)}{\partial \alpha_k} = a_k + \lambda(1 + \log \alpha_k) + u - v_k = 0, \\ v_k \alpha_k = 0, \\ \sum_k \alpha_k = 1, \alpha_k \geq 0, \\ v_k \geq 0, \end{cases}$$

for all $k = 1, 2, \ldots, K$. Assuming non-zero contributions of each modality, i.e. strictly positive values for $\alpha$, we have $v_k = 0$ for all $k$ and $\alpha_k = \exp\{(-a_i - u)/\lambda - 1\}$. Together with the constraint $\sum_k \alpha_k = 1$, we obtain

$$\alpha_k = \frac{m_k}{\sum_k m_k},$$

where $m_k = \exp\{-a_i/\lambda - 1\}$. We alternate iteratively between steps 1 and 2 until the improvement in the objective value falls below a predefined error threshold $\epsilon$ or until a maximum number *maxIter* of iterations has been reached.

#### Generalizing UMAP to multimodal data in j-UMAP

UMAP uses different definitions of high and low-dimensional similarities $p_{ij}$ and $q_{ij}$, respectively. In particular, UMAP similarities $p_{j|i}$ are defined and symmetrized to give $p_{ij}$ as follows:

$$\begin{aligned} p_{j|i} &= \exp[(-d(x_i, x_j) - \rho_i)/\sigma_i)], \qquad p_{i|i} = 0, \\ p_{ij} &= p_{j|i} + p_{i|j} - p_{j|i}p_{i|j}, \end{aligned}$$

where $\rho_i$ is the distance to the nearest neighbor of $x_i$, and $\sigma_i$ is the normalizing factor which is found through binary search using a criteria similar to the perplexity-based selection of bandwidts in t-SNE. The low-dimensional similarities are defined as:

$$q_{ij} = \left(1 + a \left\| y_i - y_j \right\|_2^{2b}\right)^{-1},$$

where $a$ and $b$ are user-defined parameters with default values $a \approx 1.929$ and $b \approx 0.7915$. Note that setting $a = b = 1$ gives the Student $t$-distribution used to define low-dimensional similarities in t-SNE (equation (3)). In contrast to t-SNE, however, $p_{ij}$ and $q_{ij}$ are not further normalized. UMAP determines the low dimensional embedding by minimizing the cross entropy:

$$C(\mathcal{E}) = \mathrm{CE}(P, Q) = \sum_{i \neq j} p_{ij} \log\left(\frac{p_{ij}}{q_{ij}}\right) + (1 - p_{ij}) \log\left(\frac{1 - p_{ij}}{1 - q_{ij}}\right), \tag{11}$$

which is equivalent to the following objective function (ignoring constant terms):

$$C(\mathcal{E}) = \sum_{i \neq j} -p_{ij} \log q_{ij} - (1 - p_{ij}) \log(1 - q_{ij}). \tag{12}$$

Objective (12) can be minimized by stochastic gradient descent.

j-UMAP, the generalization of UMAP to multimodal data is analog to j-SNE. In particular, we minimize the cross entropy between distributions $P^{(k)}$ for each modality $k$ and distribution $Q$:

$$C(\mathcal{E}) = \sum_k \alpha_k \mathrm{CE}(P^{(k)} || Q) + \lambda \sum_k \alpha_k \log \alpha_k, \tag{13}$$

where, as in the joint t-SNE objective (6), the first term denotes a convex combination with coefficients $\alpha_i$ and the second regularization term is weighted with parameter $\lambda > 0$.

Similar to j-SNE, we use an alternating optimization approach to minimize objective (13). In contrast to t-SNE, UMAP does not normalize distributions $P$ and $Q$ (compare equations (1)-(3)). To be able to set $\lambda$ to an identical value across experiments, we therefore normalize coefficients in the first term of (13) by their maximum value computed in the first iteration.

#### Supplementary Tables

Table S1: Comparison of joint embeddings and the naive concatenating approach under noise. MixRNA denotes randomly shuffled gene expression measurements between cell lines BJ and H1 while keeping scATAC-seq counts (ChrAcc) unchanged. The two rightmost columns show the results when adding random noise sampled from a uniform distribution $U[0, 100]$.

| | MixRNA+ChrAcc | | MixRNA + ChrAcc + Noise | |
|---|---|---|---|---|
| Method | KNI | ARI | KNI | ARI |
| j-SNE | 0.95 | 0.90 | 0.90 | 0.85 |
| j-UMAP | 0.95 | 0.90 | 0.94 | 0.78 |
| Concat t-SNE | 0.79 | 0.59 | 0.29 | 0.00 |
| Concat UMAP | 0.77 | 0.52 | 0.28 | 0.00 |

Table S2: Comparison of joint and unimodal embeddings on the PBMC and CBMC data sets. KNI denotes the fraction on $k$-nearest neighbors in the embedding that are of the same type, averaged over all cells. ARI measures the agreement between true cell labels and labels obtained by a Louvain clustering of the embedded points. Only cells assigned identical labels based on joint clusterings by CiteFuse and Specter are considered in the evaluation.

| | CBMC | | PBMC | |
|---|---|---|---|---|
| Method | KNI | ARI | KNI | ARI |
| j-SNE | **0.998** | **0.838** | **0.985** | **0.554** |
| RNA based t-SNE | 0.978 | 0.492 | 0.960 | 0.071 |
| ADT based t-SNE | 0.989 | 0.059 | 0.949 | 0.447 |
| j-UMAP | **0.998** | **0.927** | **0.985** | **0.682** |
| RNA based UMAP | 0.980 | 0.651 | 0.960 | 0.062 |
| ADT based UMAP | 0.982 | 0.062 | 0.948 | 0.071 |

### Supplementary Figures

#### Proof of concept

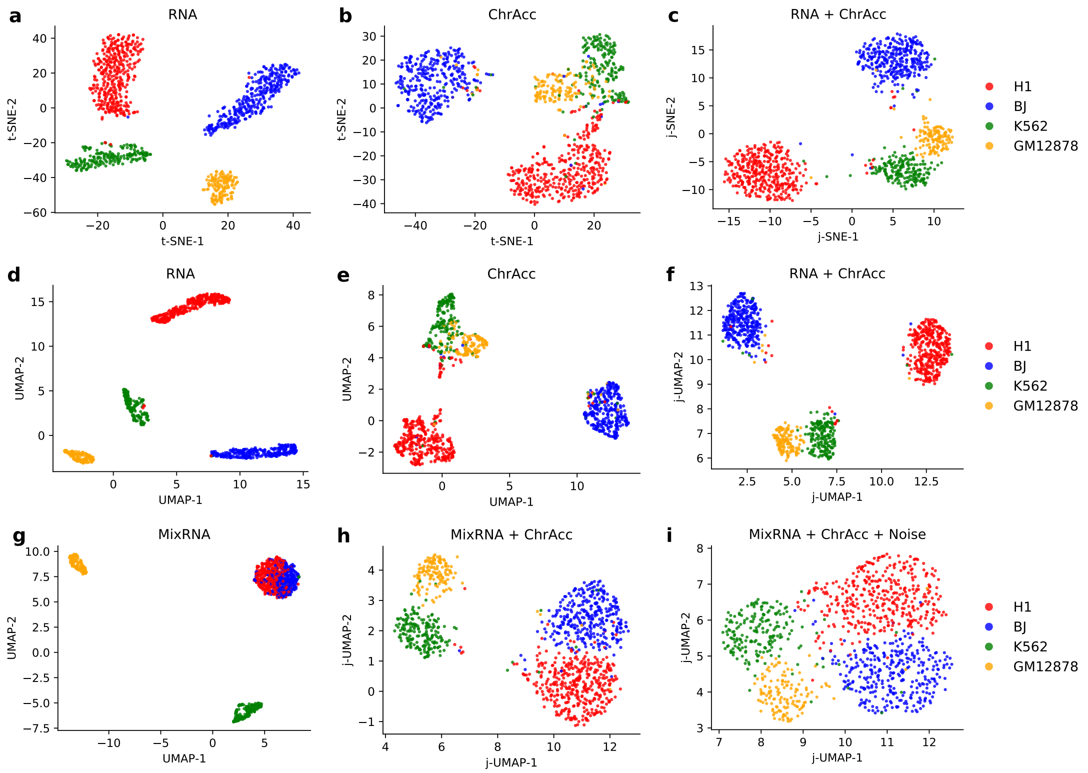

Figure S1: Unimodal and multimodal visualization of (perturbed) SNARE-seq measurements. Accessible chromatin (ChrAcc) and gene expression was measured simultaneously in single cell from human cell lines BJ, H1, K562, and GM12878. *First row:* Comparison of t-SNE and j-SNE. Conventional t-SNE embedding of measurements of (**a**) RNA or (**b**) accessible chromatin. (**c**) Joint embedding of both modalities (RNA and chromatin accessibility) by j-SNE. *Second row:* Comparison of UMAP and j-UMAP. Conventional UMAP of (**d**) RNA or (**e**) accessible chromatin. (**f**) Joint embedding of both modalities by j-UMAP. *Third row:* RNA measurements were randomly shuffled between cell lines BJ and H1 (MixRNA). (**g**) Conventional UMAP embedding of cells based on shuffled gene expression alone. (**h**) j-UMAP visualization of shuffled gene expression and (unchanged) chromatin accessibility. (**i**) j-UMAP embedding of shuffled gene expression, chromatin accessibility, and a third modality that consist entirely of random noise sampled from uniform distribution $U[0, 100]$. The simultaneously optimized weights assigned to the three modalities by JVis were $\alpha = (0.32, 0.43, 0.25)$, effectively suppressing noise.

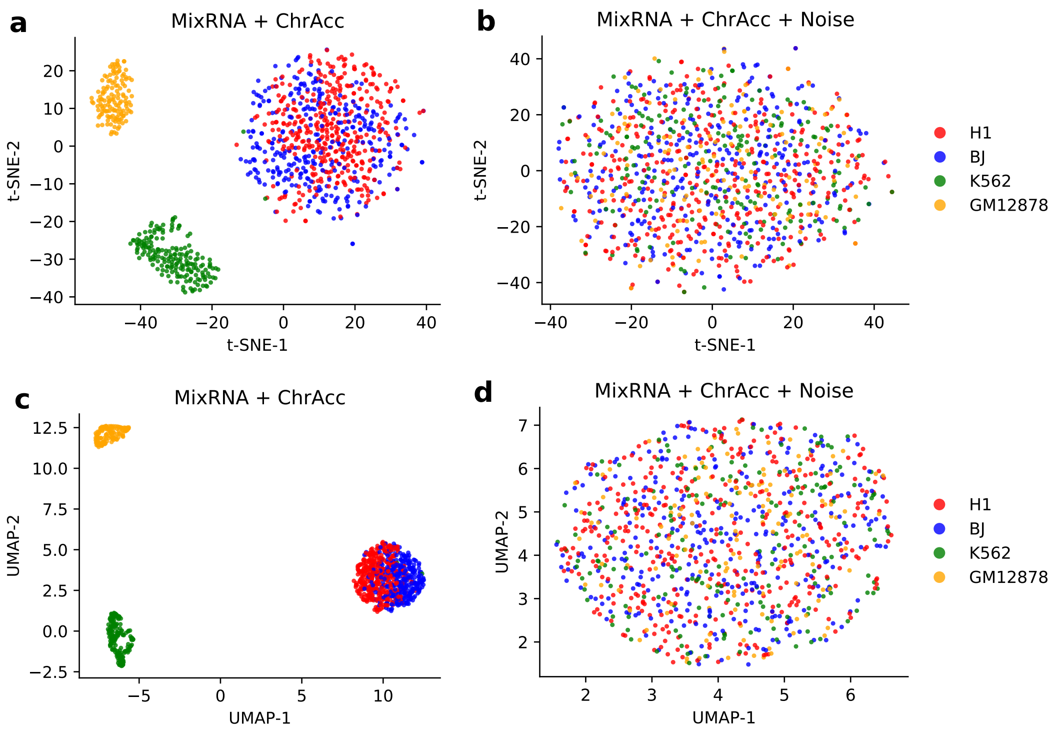

Figure S2: Visualization of concatenated vectors for four cell lines with perturbation. Conventional t-SNE (**a**, **b**) and UMAP (**c**, **d**) embeddings of concatenated vectors containing shuffled gene expression measurements between cell lines BJ and H1 and (unchanged) chromatin counts (**a**, **c**). (**b**, **d**) uniform noise was added as third modalitiy.

#### JVis agrees with joint clustering

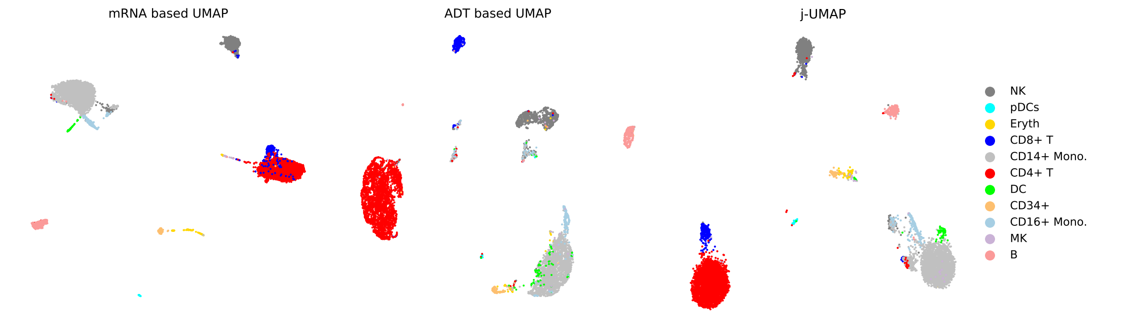

Figure S3: UMAP/j-UMAP visualization of CBM cells. Cluster labels were identified by Specter. Embeddings were computed from mRNA measurements alone (left), protein expression (ADT) alone (middle), or jointly from both (right).

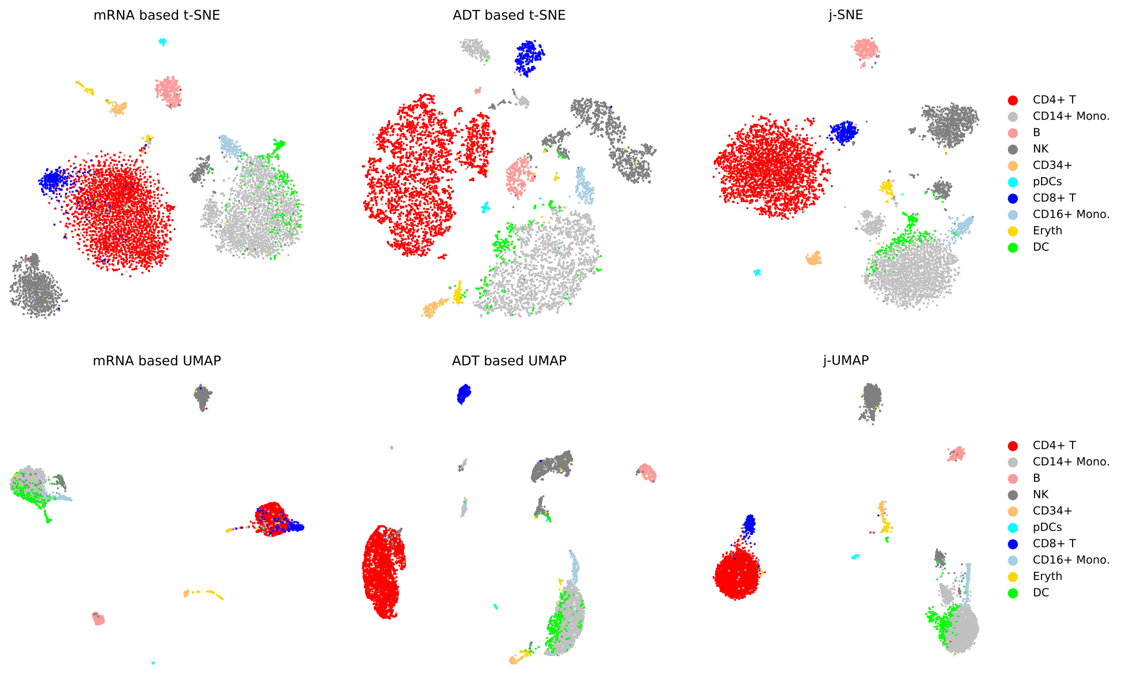

Figure S4: t-SNE/j-SNE (top row) and UMAP/j-UMAP (bottom row) visualizations of CBM cells. Cluster labels were identified by CiteFuse. Embeddings were computed from mRNA measurements alone (left), protein expression (ADT) alone (middle), or jointly from both (right).

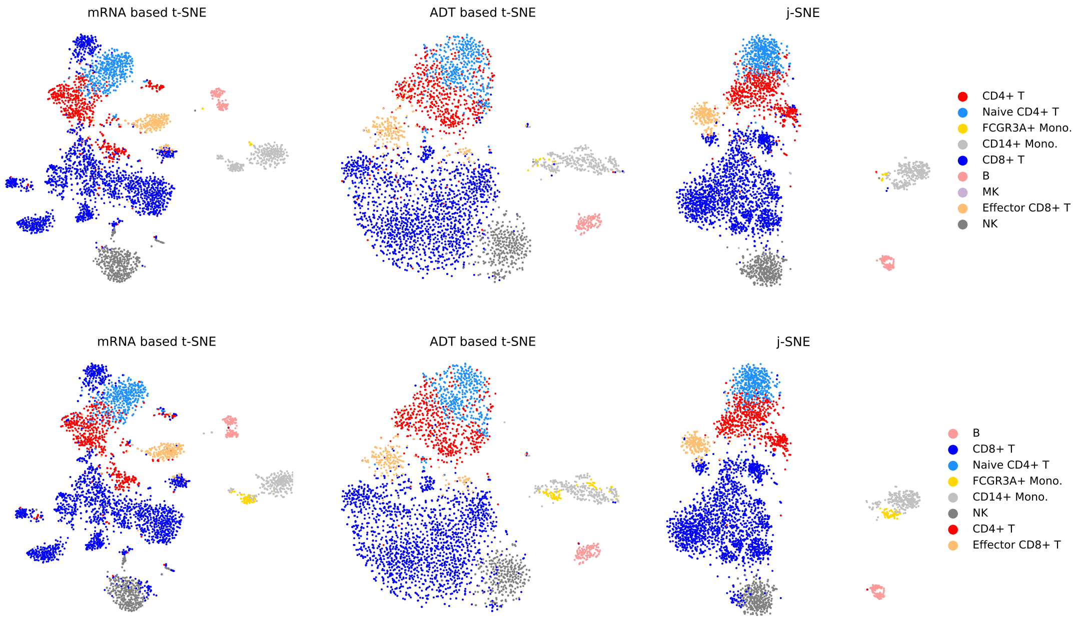

Figure S5: t-SNE/j-SNE visualizations of PBM cells. Cluster labels were identified by Specter (top row) or CiteFuse (bottom row). Embeddings were computed from RNA measurements alone (left), protein expression (ADT) alone (middle), or jointly from both (right).

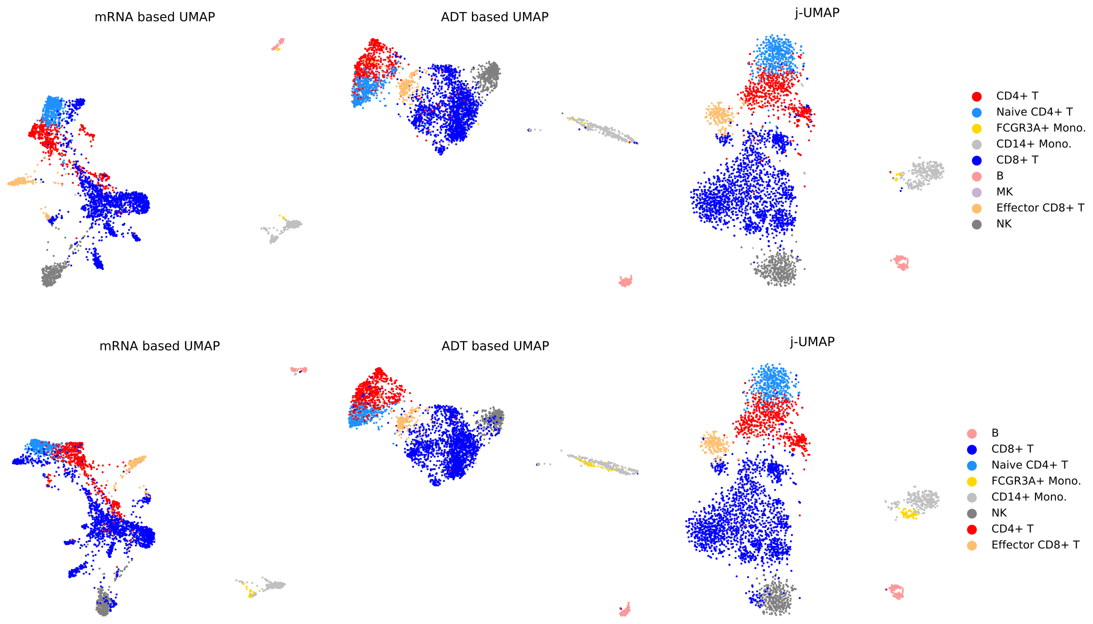

Figure S6: UMAP/j-UMAP visualization of PBM cells. Cluster labels were identified by Specter (top row) or CiteFuse (bottom row). Embeddings were computed from RNA measurements alone (left), protein expression (ADT) alone (middle), or jointly from both (right).

#### Protein acceleration in joint embeddings

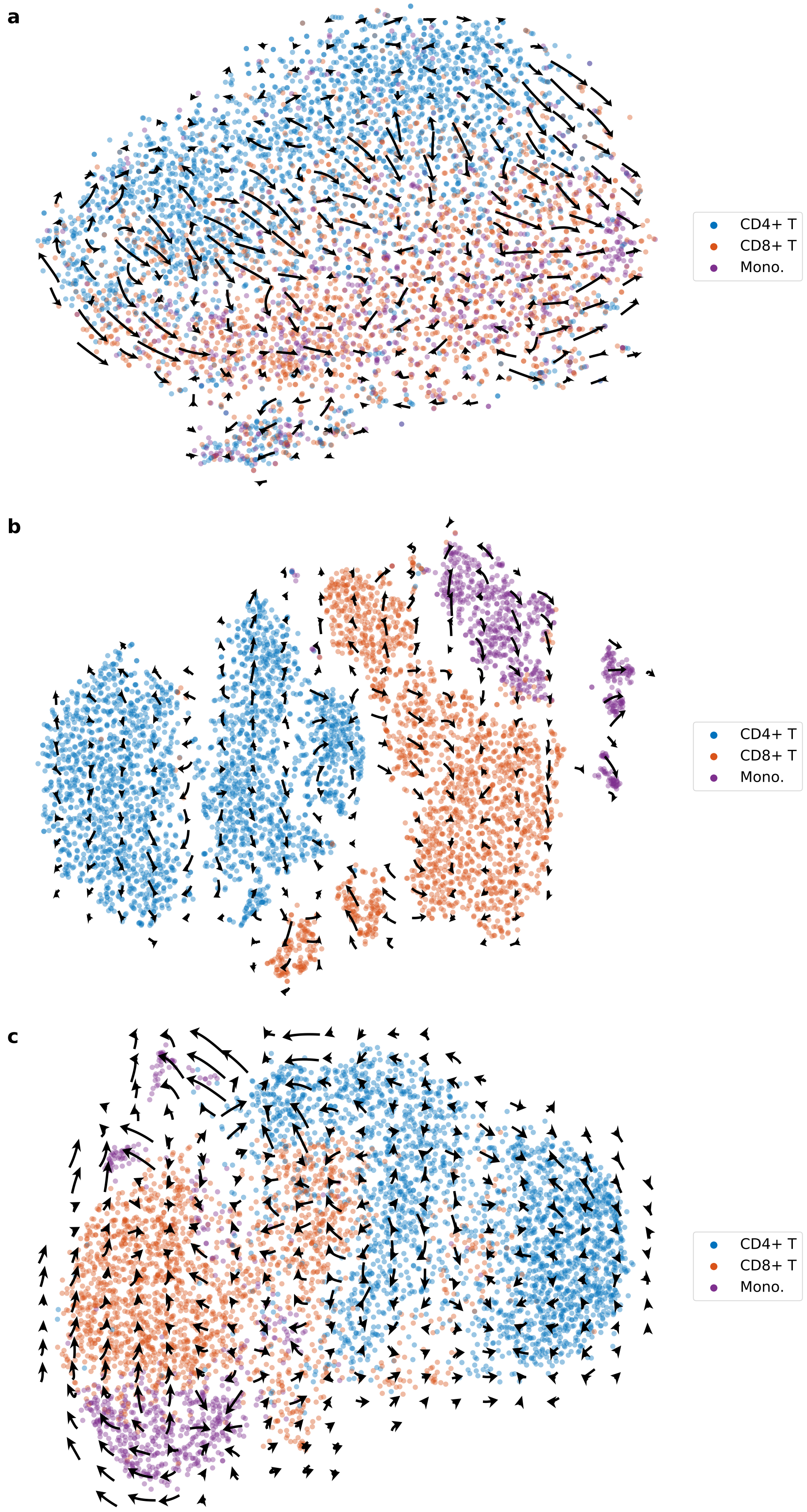

Figure S7: Protein acceleration in ECCITE-seq data set CTCL projected into transcriptome-based t-SNE (a), j-SNE (b), and j-UMAP (c) embeddings.

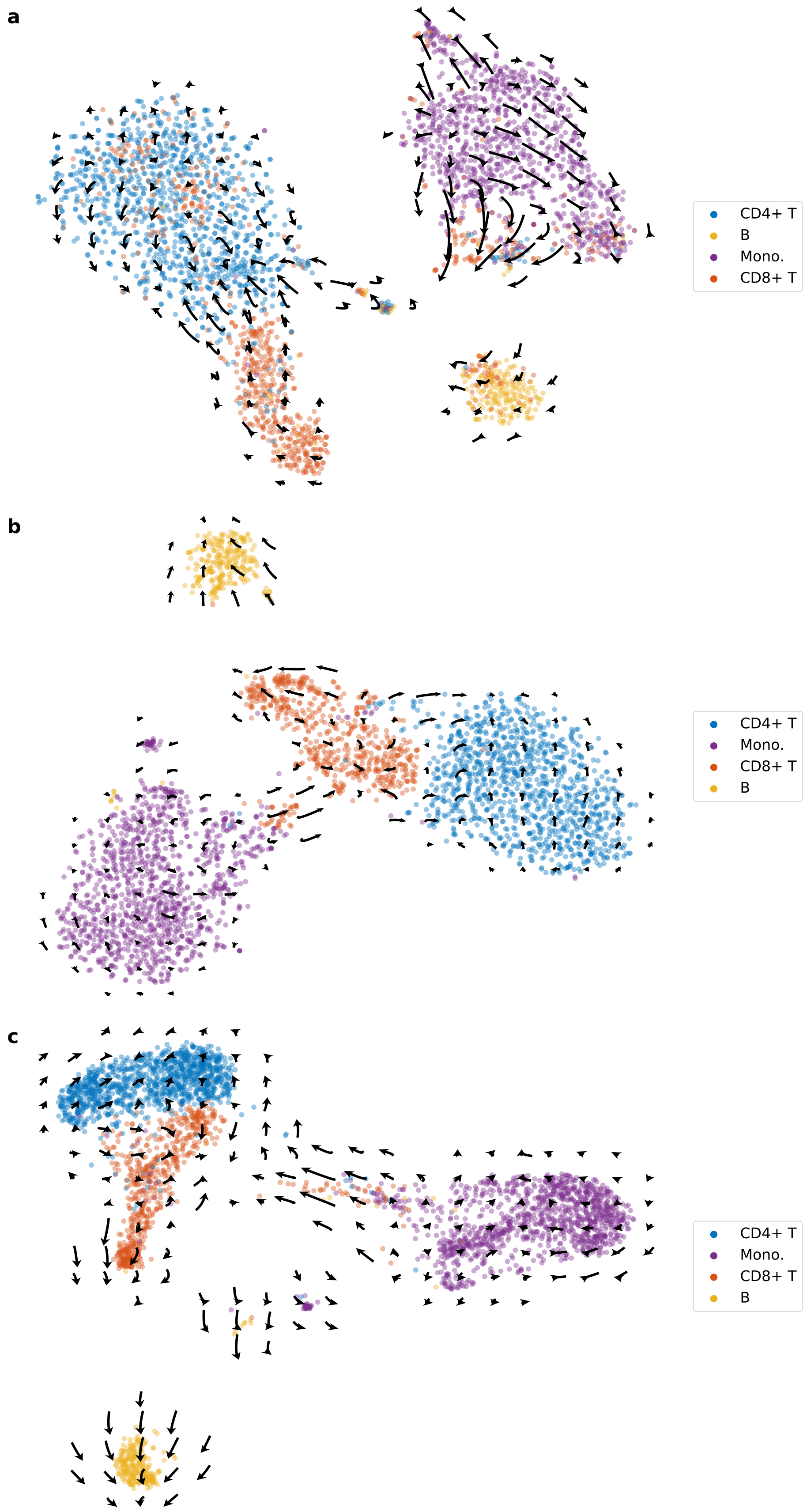

Figure S8: Protein acceleration in REAP-seq data set projected into transcriptome-based t-SNE (a), j-SNE (b), and j-UMAP (c) embeddings.

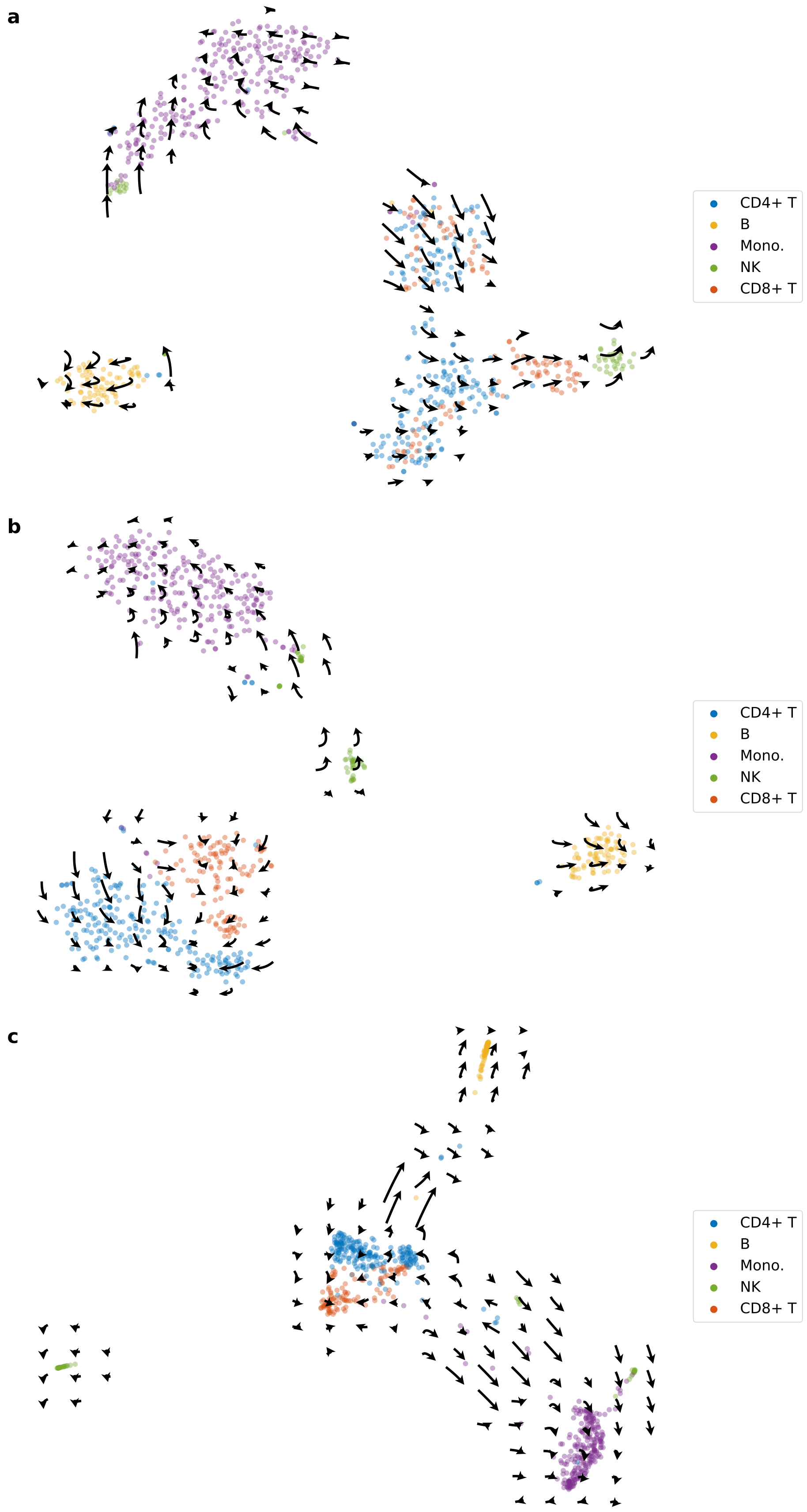

Figure S9: Protein acceleration in 10X 1k data set projected into transcriptome-based t-SNE (a), j-SNE (b), and j-UMAP (c) embeddings.

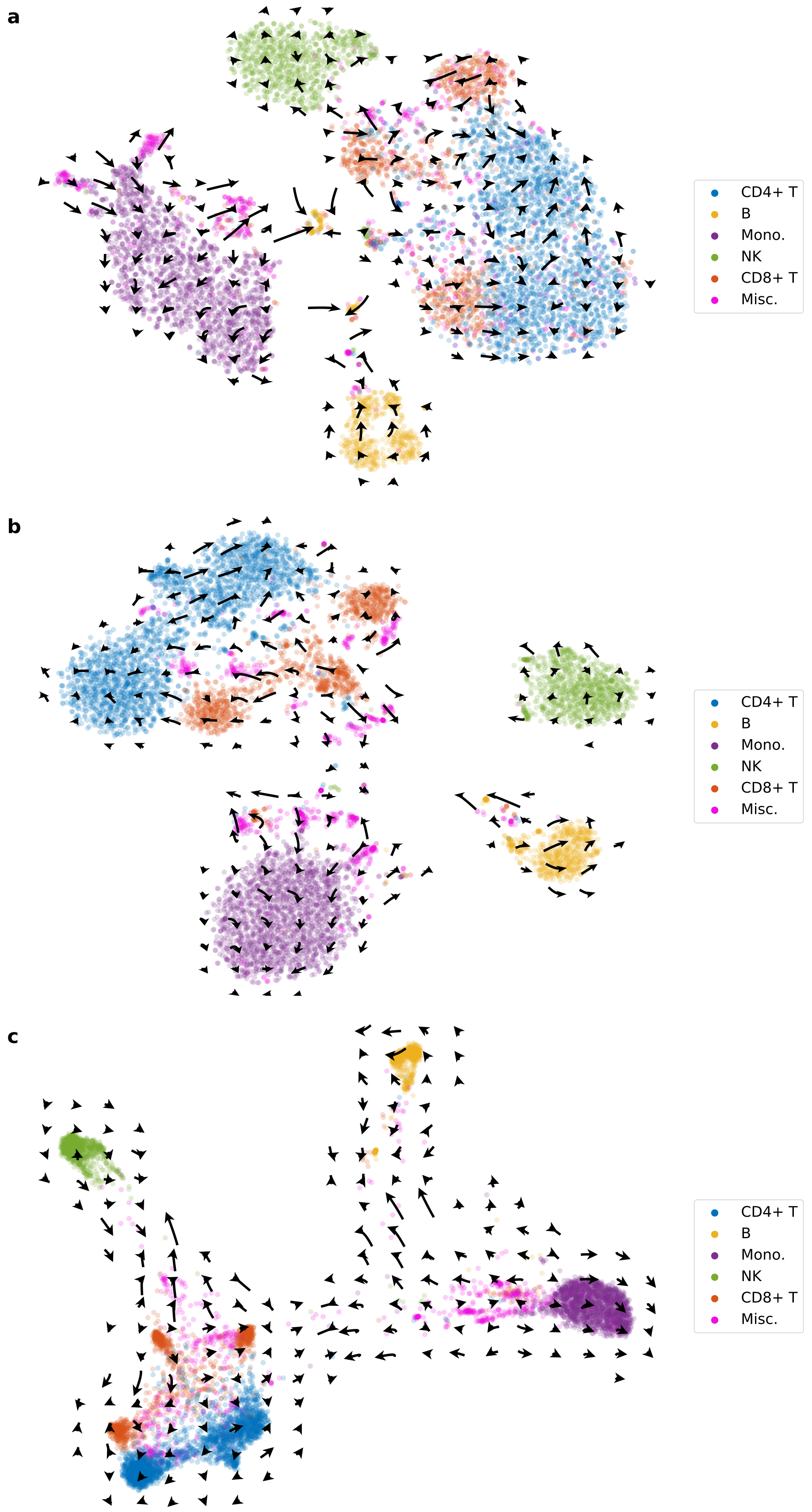

Figure S10: Protein acceleration in 10X 10k data set projected into transcriptome-based t-SNE (a), j-SNE (b), and j-UMAP (c) embeddings.

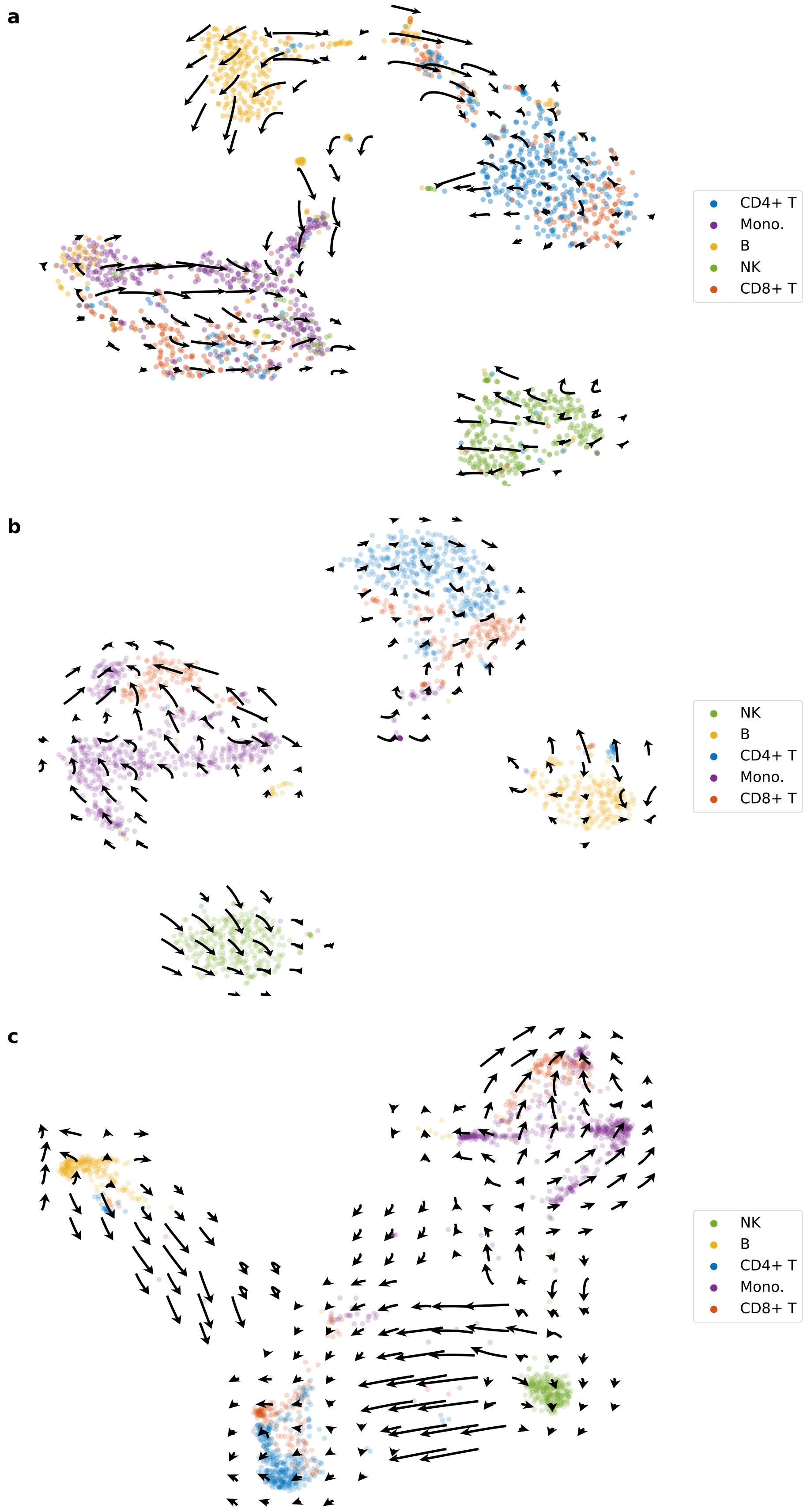

Figure S11: Protein acceleration in CITE-seq data set projected into transcriptome-based t-SNE (a), j-SNE (b), and j-UMAP (c) embeddings.

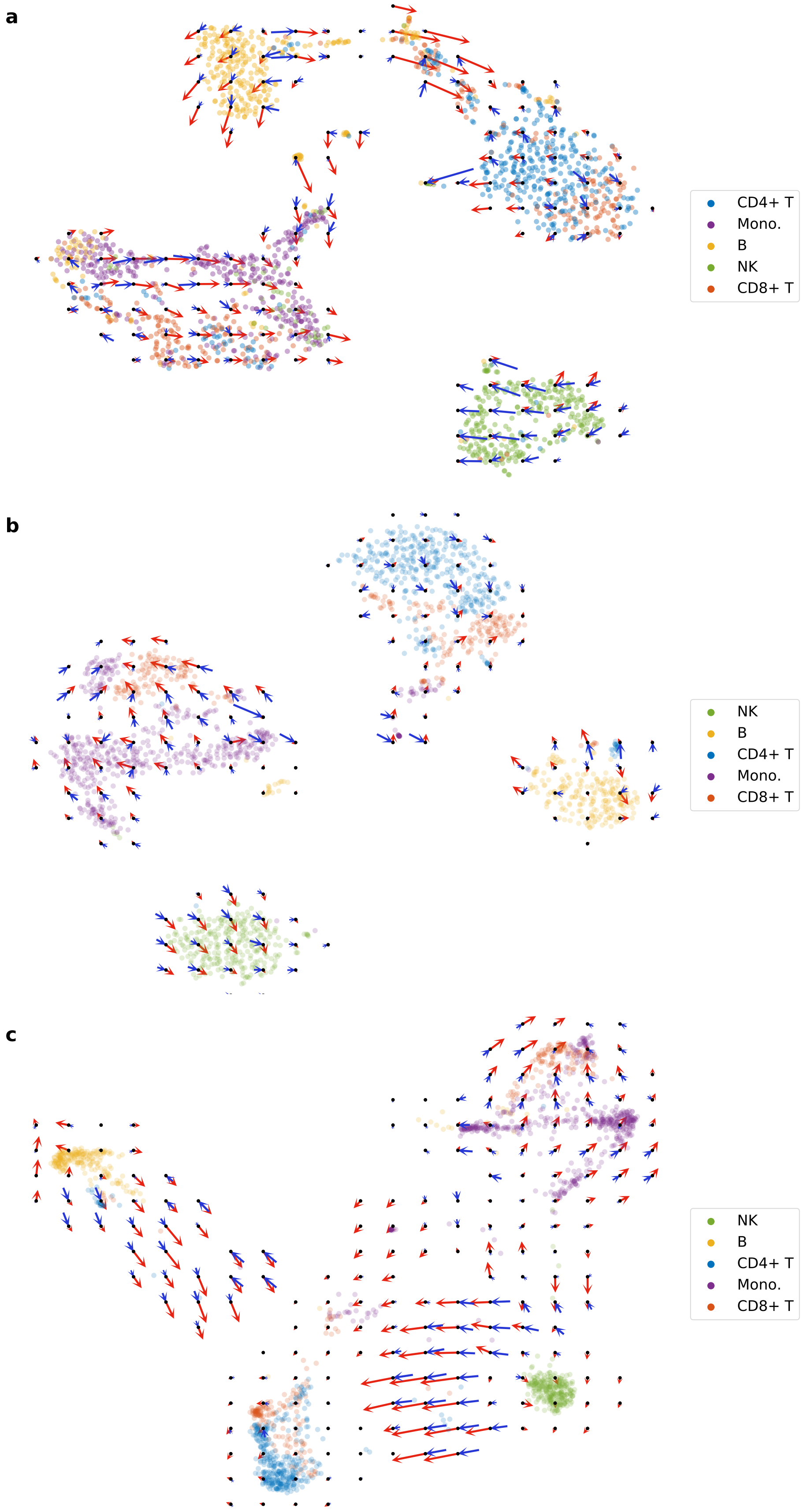

Figure S12: CITE-seq velocity landscapes. RNA velocities (red) and protein velocities (blue) are projected into transcriptome-based t-SNE (a), j-SNE (b), and j-UMAP (c) embeddings.

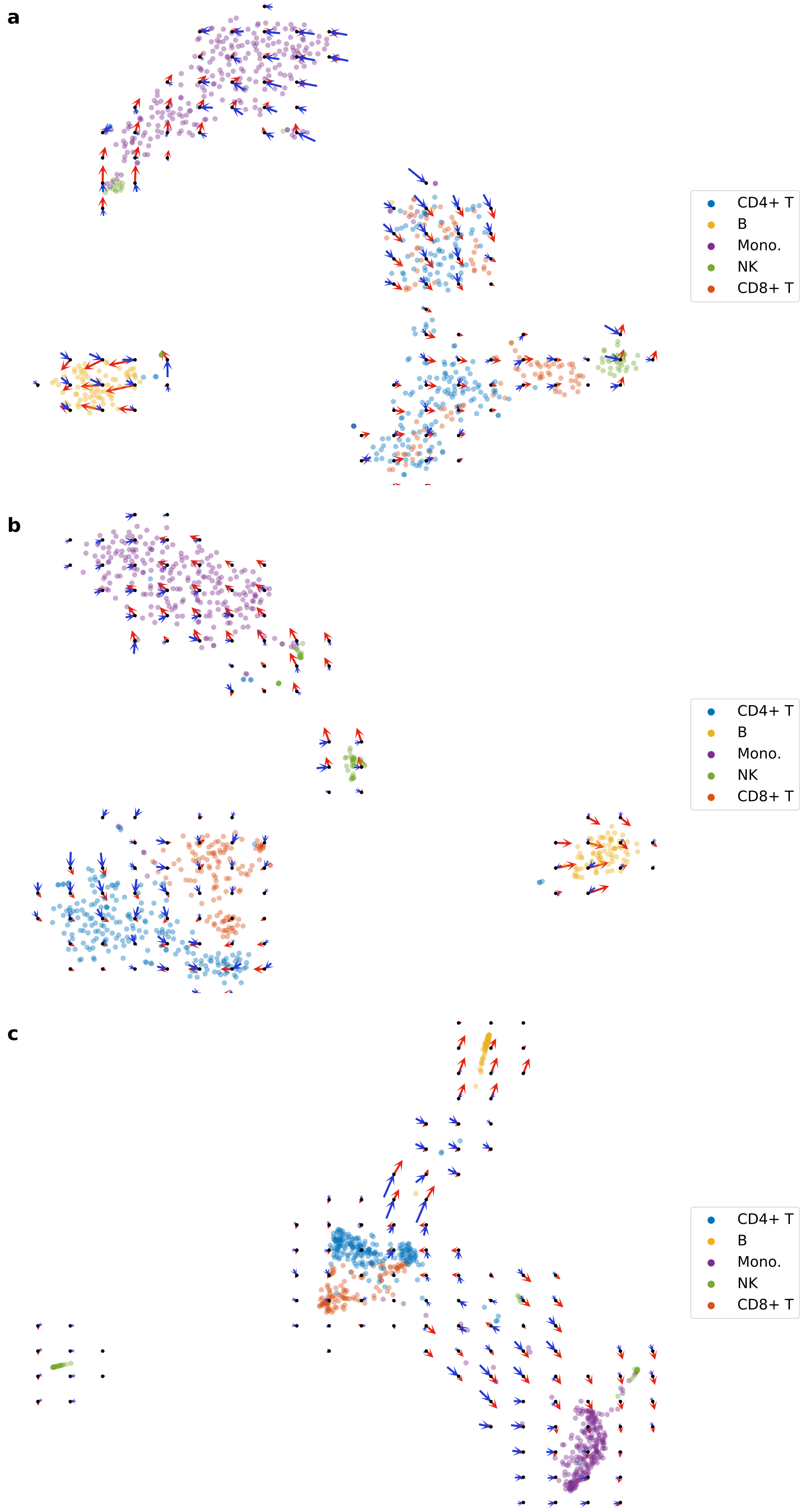

Figure S13: 10X 1k velocity landscapes. RNA velocities (red) and protein velocities (blue) are projected into transcriptome-based t-SNE (a), j-SNE (b), and j-UMAP (c) embeddings.

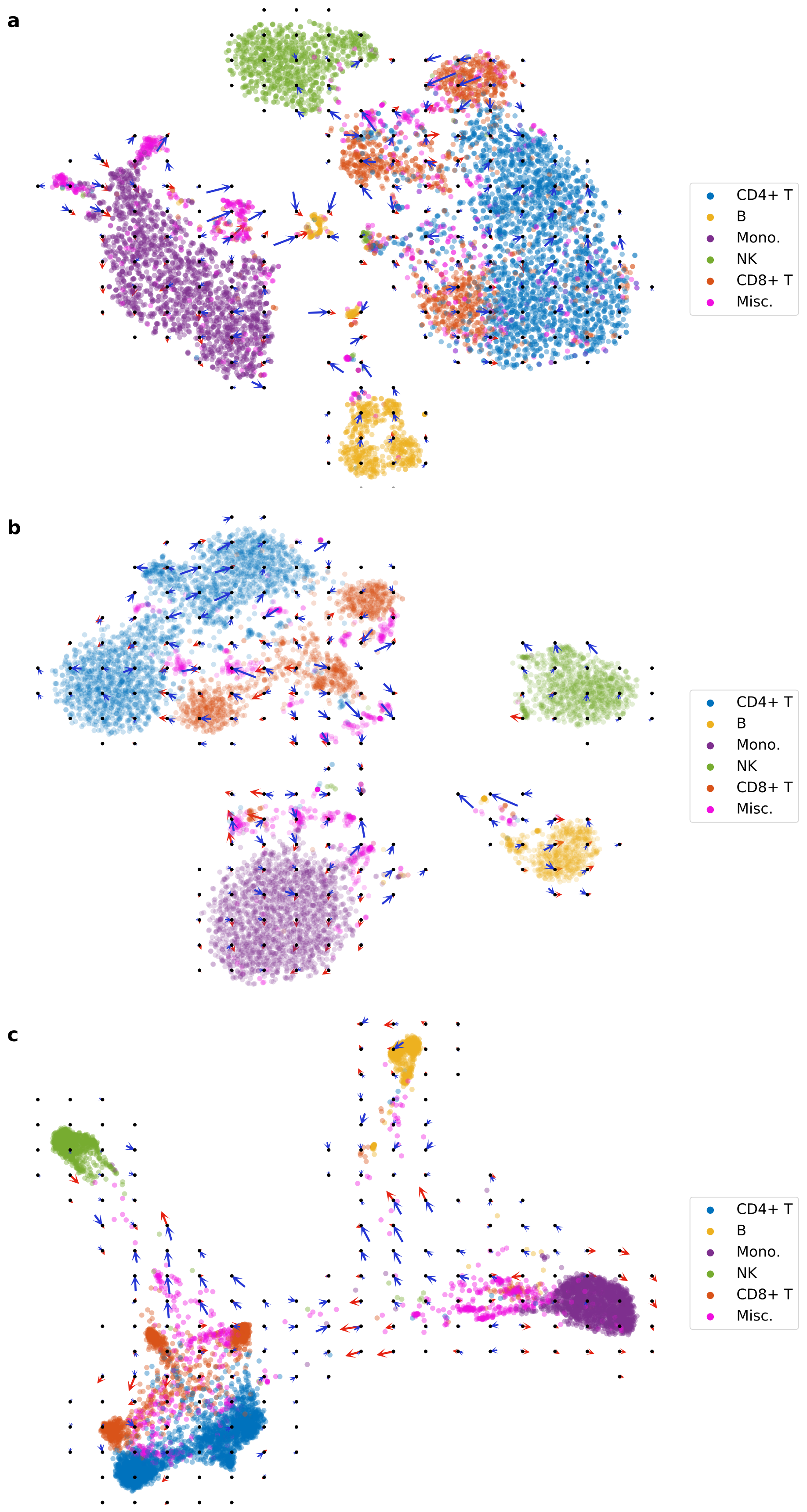

Figure S14: 10X 10k velocity landscapes. RNA velocities (red) and protein velocities (blue) are projected into transcriptome-based t-SNE (a), j-SNE (b), and j-UMAP (c) embeddings.

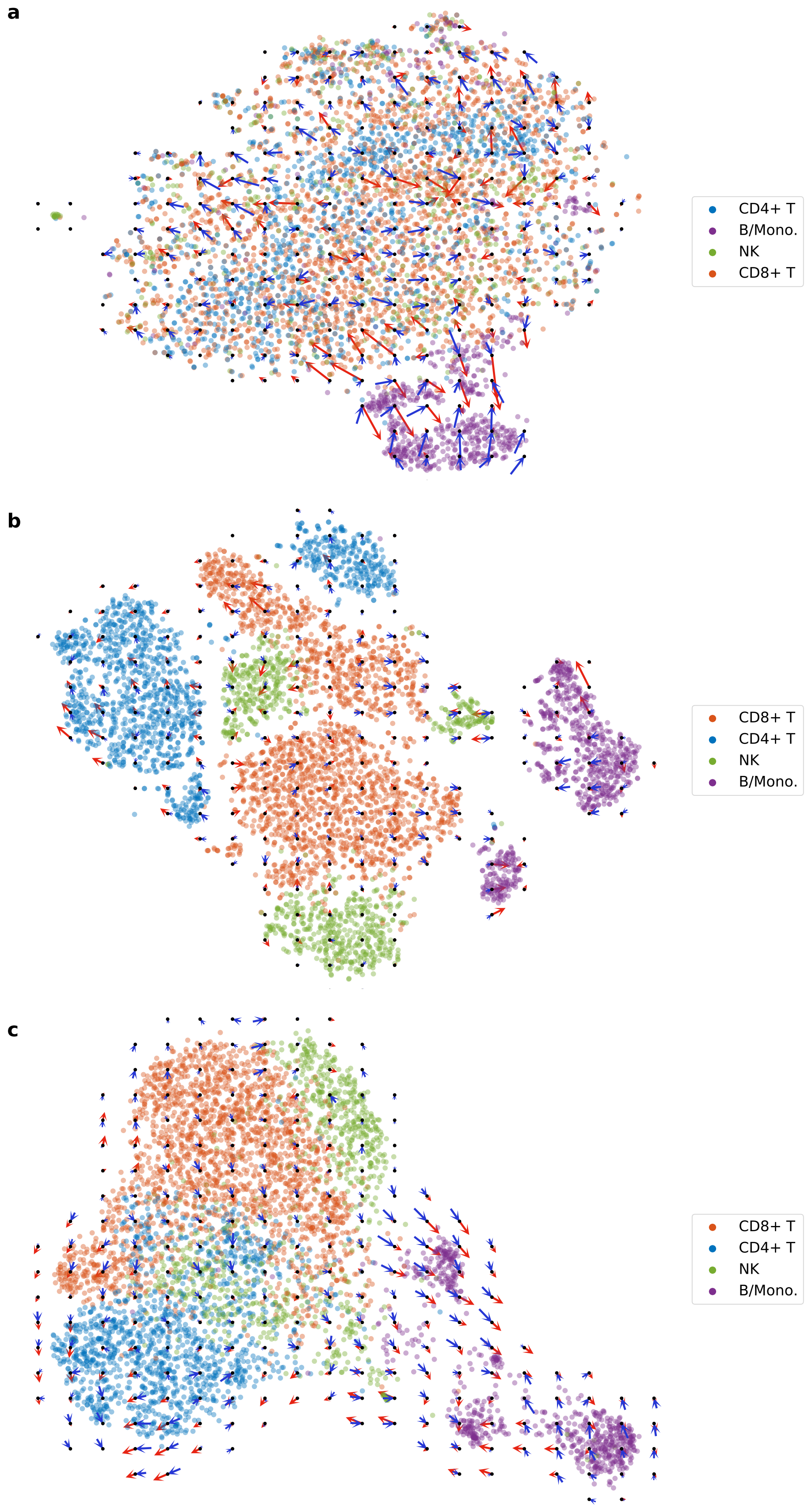

Figure S15: ECCITE-seq (ctrl) velocity landscapes. RNA velocities (red) and protein velocities (blue) are projected into transcriptome-based t-SNE (a), j-SNE (b), and j-UMAP (c) embeddings.

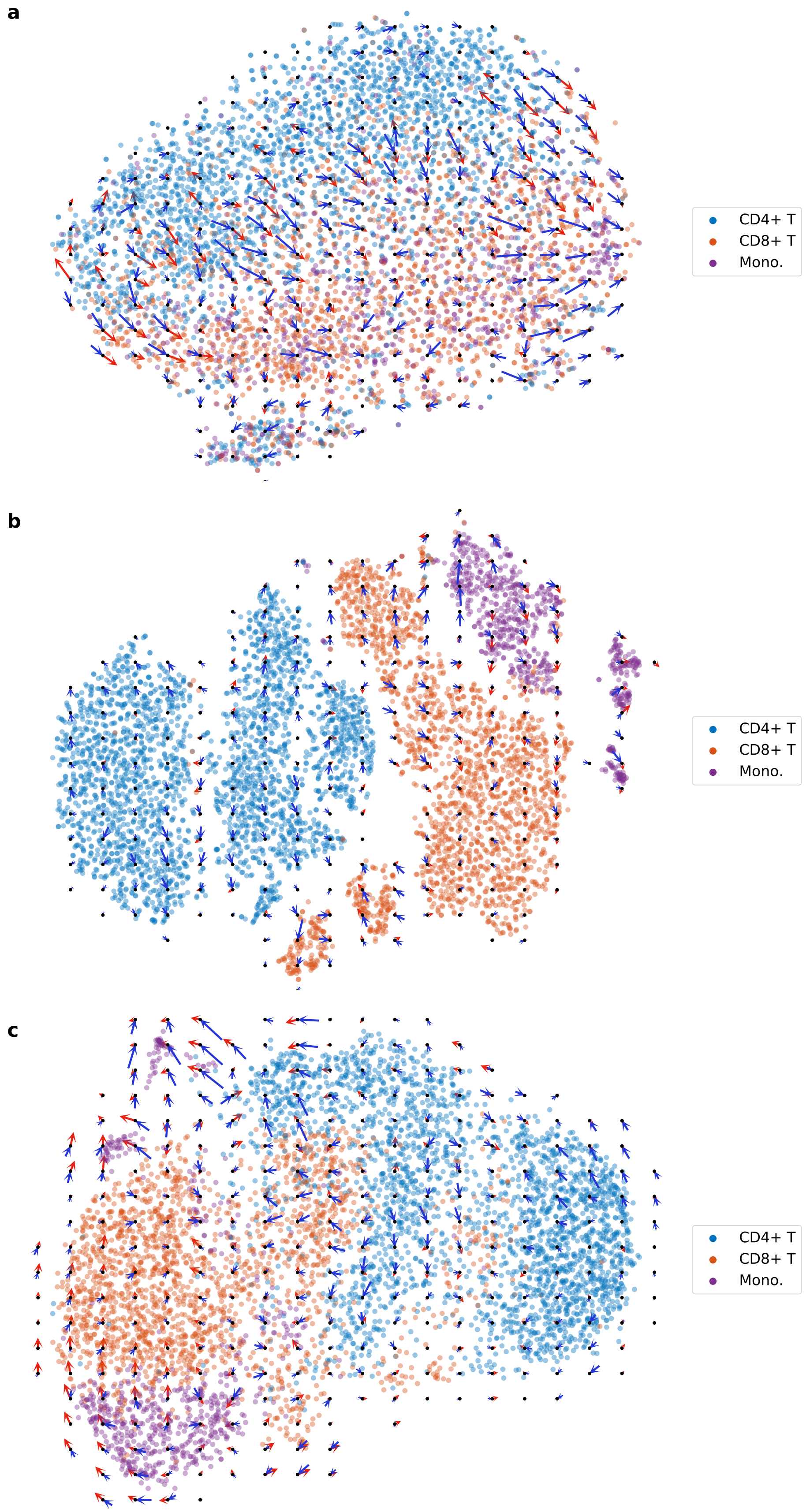

Figure S16: ECCITE-seq (CTCL) velocity landscapes. RNA velocities (red) and protein velocities (blue) are projected into transcriptome-based t-SNE (a), j-SNE (b), and j-UMAP (c) embeddings.

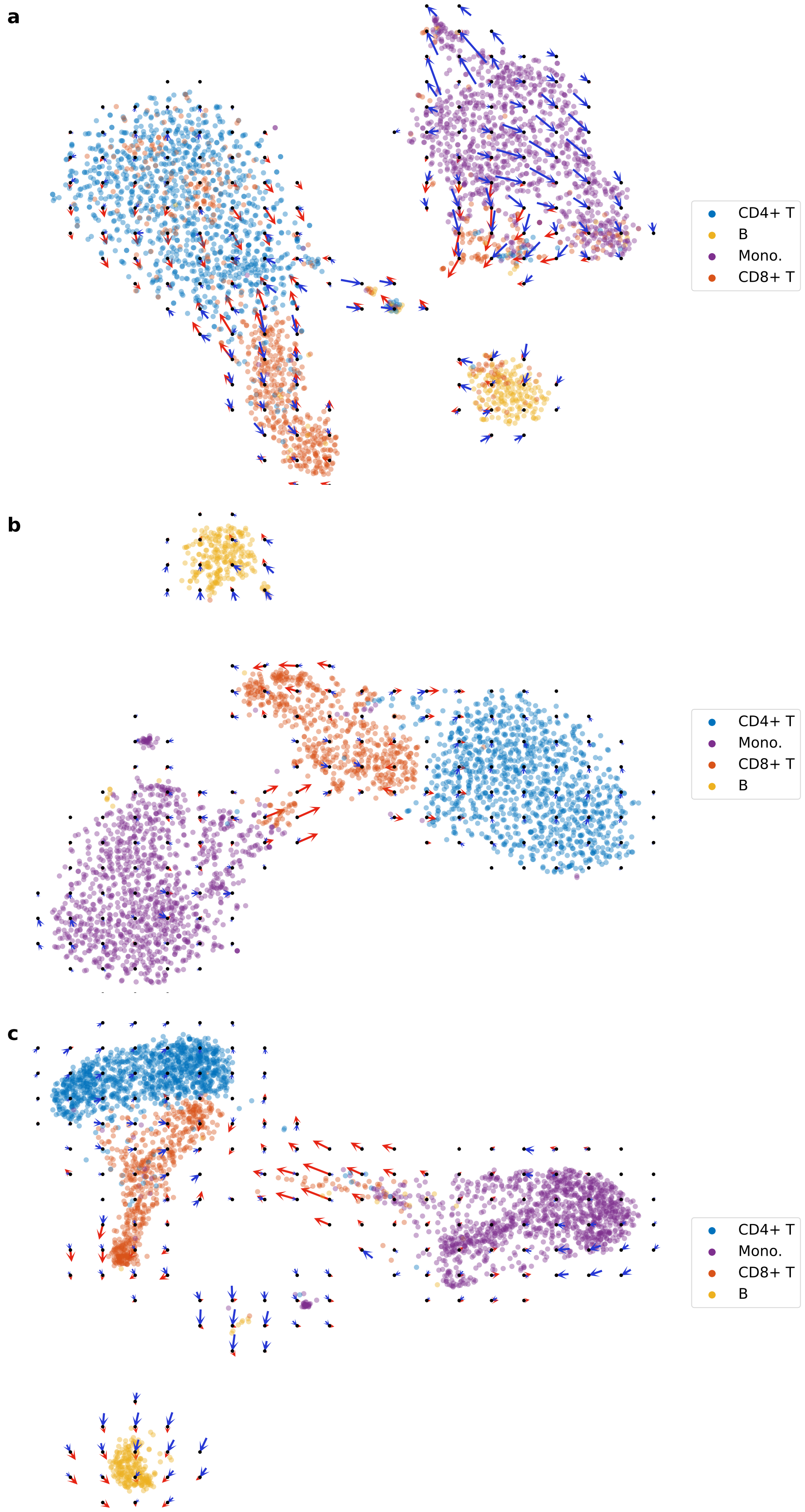

Figure S17: REAP-seq velocity landscapes. RNA velocities (red) and protein velocities (blue) are projected into transcriptome-based t-SNE (a), j-SNE (b), and j-UMAP (c) embeddings.
